# Chlorotoxin does not target matrix metalloproteinase-2 in glioblastoma

**DOI:** 10.1101/2025.07.11.664294

**Authors:** Eli Blaney, Meron Demeke, Seraphine Kamayirese, Louise Monga, Laura A. Hansen, Charles R. Watts, Sándor Lovas

**Affiliations:** Department of Biomedical Sciences, School of Medicine, Creighton University, Omaha, NE 68178; Department of Neurosurgery, Park Nicollet, Methodist Hospital, St. Louis Park, MN 55426

## Abstract

Glioblastoma aggressively invades surrounding tissue by expressing matrix metalloproteinase-2 (MMP-2). Therefore, effective inhibition of MMP-2 is a desirable target for treatment. In some reports, the chlorotoxin (Ctx) polypeptide produced by the scorpion *Leiurus quinquestriatus,* interacts with human MMP-2 to inhibit tumor invasion without affecting surrounding tissue. We employed three molecular docking methodologies followed by molecular dynamics simulations to find consensus binding and calculate the binding energy of these peptide ligands to MMP-2. In addition to the Ctx itself, four C-terminal fragments were chosen to study their binding to MMP-2. The molecular docking platforms HPEPDOCK, HADDOCK, and AlphaFold2 created peptide – protein poses for each candidate binding to MMP-2. These poses underwent 500 ns molecular dynamics simulations. Peptide binding on MMP-2 and final binding energies were calculated using the Molecular Mechanics Poisson-Boltzmann Surface Area method. Configurational entropy and root-mean square deviation analyses showed stable peptide – protein complexes. Ctx and its peptide fragments frequently bound to regions on MMP-2 other than the catalytic site. All docking methods shared consensus on large negative binding energies, indicating favorable interaction between Ctx and its analogs with MMP-2. While Ctx and its fragments bind to MMP-2, there is no consensus on which region of MMP-2 they are bound to or which peptide binds strongest. Neither Ctx nor its fragments inhibited MMP-2 enzymatic activity, however, glioblastoma cellular migration was inhibited. Interactions with the non-catalytic regions of MMP-2 suggest allosteric binding to MMP-2. Inhibition of cellular migration without inhibition of MMP-2 activity warrants further study into the possible targets of Ctx expressed in glioblastoma.

## Introduction

Glioblastoma multiforme (GBM) is a near universally fatal central nervous system tumor arising from the astrocyte glial cells [1]. The invasion of GBM cells into the extracellular matrix is mediated to a major extent by the proteolytic activity of human matrix metalloproteinase enzymes (MMPs) on basement membrane collagens [2]. In humans, the 24 isoforms of MMPs [3] are widely distributed in different organs. MMPs are metalloenzymes, containing Zn2+ ions in their active site. They share a common structure including: a propeptide, a catalytic metalloproteinase domain, a linker peptide, and a hemopexin domain [4]. The activity of MMPs is low or negligible in healthy tissues and upregulated by inflammatory cytokines, growth factors, hormones, cell − cell and cell – extracellular matrix interactions [5]. Their activity can be downregulated by tissue inhibitors of metalloproteinases (TIMPs) [6]. The delicate control of MMP expression and activity maintains the stability of tissue(s) and appropriate repair and remodeling. Abnormal tissue turnover by MMPs is leveraged by nervous system tumors to facilitate tumorigenesis [7]. Because of the high structural similarities between MMPs in normal and malignant tissue, inhibitors of MMPs have not resulted in tangible benefits [8], and development of specific inhibitors are impeded by the difficulty of targeting tumor cells while leaving normal tissue undamaged.

GBM cell lines have been shown to overexpress both MMP–2 and MMP–9 which may allow for efficient breakdown of the extracellular matrix (ECM) and subsequent migration [9]. MMP-2 comprises of a prodomain, collagenase-1 domain, collagen binding domain containing three repeats of a fibronectin type-II domain, collagenase-2 domain, and a hemopexin domain (Fig 1 and Table 1) [5]. The hinge region links these domains together. Proteolytic cleavage of prodomain results in the mature and catalytically active 62 kDa protease [6]. Previous investigations into the interaction energies of MMP-2 revealed that peptides binding the metal ions of MMP-2 decrease system entropy, contributing with residue interactions to create a large hemopexin domain movement [10]. The role of MMP-2 in angiogenesis [11] and tumor migration and invasion [12,13] contributes to the poor prognosis of patients with GBM [12,14,15]. MMP-2 has also been linked to similar detrimental effects in lung cancers [16]. Inhibition of MMP-2 has decreased GBM cell migration in cell cultures [17]. The design of selective inhibitors has however been elusive secondary to the highly conserved nature of the catalytic site across multiple expressed metalloproteinases [18].

**Fig 1.**
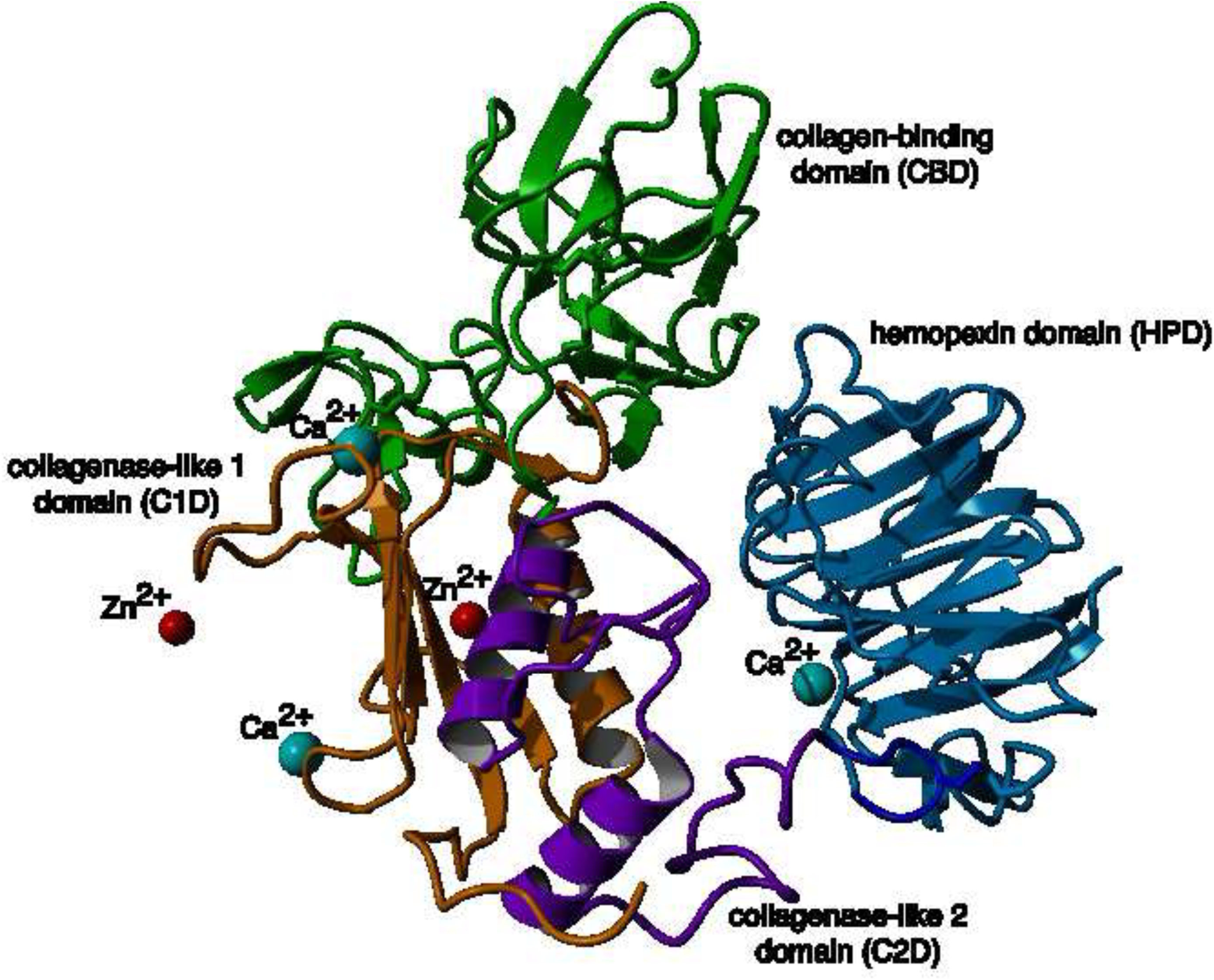
Ribbon representation of the structure of MMP-2 (PDB 1CK7). Domains identified as follows: collagenase-like 1, orange; collagen binding, green; collagenase-like 2, purple; hinge region, gray; hemopexin, red; Zn2+, red; Ca2+, cyan.

**Table 1.** Regions of matrix metalloproteinase-2.

| Amino acid residue | Region* |
| --- | --- |
| 1-109 | Prodomain |
| 110-221 | Collagenase-like 1 domain (C1D) |
| 222-396 | Collagen-binding domain (CBD) |
| 228-276 | Fibronectin type-II repeat 1 |
| 286-334 | Fibronectin type-II repeat 2 |
| 344-392 | Fibronectin type-II repeat 3 |
| 397-465 | Collagenase-like 2 domain (C2D) |
| 472-660 | Hemopexin domain (HPD) |
| 472-516 | Hemopexin repeat 1 |
| 517-563 | Hemopexin repeat 2 |
| 565-613 | Hemopexin repeat 3 |
| 614-660 | Hemopexin repeat 4 |

\*C1D, C2D, C2D and HPD abbreviations are in Fig.1

One such proposed binder to MMP-2 includes Chlorotoxin (Ctx). Ctx (Fig 2) is a 36-residue polypeptide from the scorpion *Leiurus quinquestriatus* [19,20]. Ctx contains 8 cysteinyl residues, forming 4 disulfide bridges which help to stabilize its three-dimensional structure [21,22]. Initially, it was shown to inhibit small conductance chloride channels on the surface GBM cells and other tumors of neuroectodermal embryologic origin as well as possible inhibition of MMP-2 [19,23–25]. However, conflicting data on the effects of Ctx on the glioma-specific chloride channel makes it difficult to confirm if Ctx targets chloride channels at a high affinity [25–27]. Deshane and colleagues showed the overexpression of MMP-2 on the surface of GBM cells and the binding of Ctx to MMP-2 [28]. They also determined that Ctx inhibits migration of GBM cells with an IC50 of 184 nM and inhibits MMP-2 enzymatic activity [28]. Ctx has emerged as a promising target for therapeutic intervention because it selectively recognizes and binds to GBM cells while sparing normal brain cells [29]. This selectivity was confirmed in a previous study which found that Ctx binds to tumors of neuroectodermal origin and is unable to bind to non-tumorigenic neurological samples [25,30–32]. Although an X-ray crystallographic structure of Ctx bound to MMP-2 has not been published, molecular dynamics simulations have shown a possible interaction between the C-terminal residues of Ctx and the collagen binding domain of MMP-2 [33].

**Fig 2.**
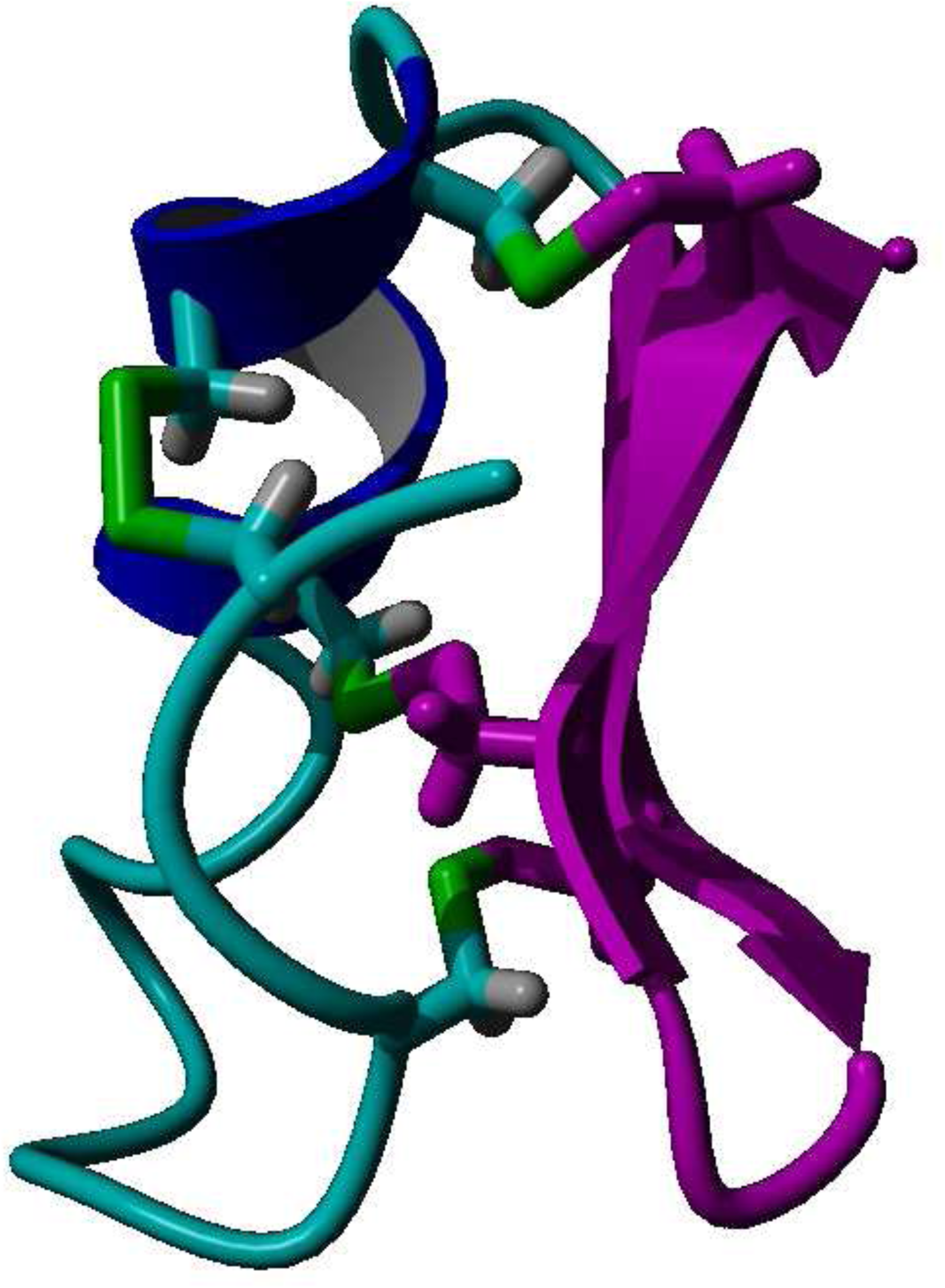
Ribbon representation of the three-dimensional structure of Ctx. (**A**) Random coil, cyan; alpha helix, blue; beta turn, green; residues Gly 24 – Arg 36, pink. Disulfide bridges are shown in sticks. (**B**) amino acid sequence of Ctx. Disulfide bridges are present between residue pairs (2,19), (5,28), (16,33), (20,35).

Although MMP-2 has been identified as a potential target protein of Ctx, its interaction and impact on MMP-2 enzymatic activity remain to be fully elucidated. While some publications claim that Ctx can inhibit the gelatinase activity of MMP-2 [28,34], Farkas and colleagues were unable to show any significant biological effect of Ctx on the activity of MMP-2 [35]. They also showed that Neuropilin-1 (NRP-1) binds both the free carboxyl and amide protected C-terminal forms of Ctx. This conflicting data prompted us to investigate whether Ctx can bind to and suppress MMP-2 activity.

Even though the inhibitory effects of Ctx on GBM cells remains elusive with regards to etiology, Ctx remains clinically relevant for the potential management of GBM with three registered clinical trials within the United States [36–38]. The K23R mutated Ctx conjugated to indocyanine green (tozuleristide, BLZ-100) is in clinical trials as an intraoperative fluorescent tumor dye to assist with improved resection in both GBM and skin cancers [36–39]. Tozuleristide/BLZ-100 also binds to the vascular epithelium of cerebral vascular malformations (cavernous vascular malformations) indicating another possible use for this agent in surgical practice [40]. Ctx is also used in a chimeric antigen receptor T-cell (CAR-T) immunotherapy strategy targeting GBM [41,42]. The CAR-T construct consists of Ctx, an IgG4Tc spacer, and a CD28 co-stimulatory domain expressed on engineered and expanded CD19 cell lines. These therapies have shown considerable efficacy in the treatment of hematopoietic malignancies [43]. Given the promise of these potential therapeutics, further characterization of Ctx binding to GBM cells is warranted.

Othman *et al.* using molecular docking and molecular dynamics (MD) simulations showed that Ctx binds to MMP-2 at its collagen binding domain [33]. This data suggests that the C-terminal region of Ctx interacts with MMP-2. Dastpeyman *et al.*, using synthetic fragments of Ctx, showed that the last eight-residue fragment of Ctx inhibits GBM cell migration [44]. Using extensive MD simulations, we have shown that substituting the C-terminal Cys residues with either Ser or α-amino-butyric acid (Abu) residues the β-hairpin conformation is retained [22].

To determine if Ctx binds and inhibits MMP-2, we also designed C-terminal fragments that encompass residues used by Dastpeyman [44] and expanded upon the computational approaches of Othman [33], employing three molecular docking methodologies followed by MD simulations. Molecular docking provides only poses of binding; therefore, they must be followed by an additional investigation to describe the detailed interactions between the ligands and receptor. These four methods in total were compared to find consensus binding and calculate the binding energies of these peptide ligands to MMP-2. This computational approach was complemented by differential scanning fluorimetry (DSF) and surface plasmon resonance (SPR) assays. Inhibition of MMP-2 enzymatic activity and inhibition of its glioblastoma cell migration by peptides were also assessed through an MMP-2 inhibition assay and a wound-healing scratch assay.

## Materials and Methods

### Peptides and Proteins

C-terminally amide protected Ctx, and peptide analogs were obtained from Biosynth International Inc (Louisville KY, USA). All peptide analogs were N- and C-terminally acetyl and amide protected, respectively. Peptides were greater than 95% pure. Recombinant MMP-2 (UniProt: P08253) was obtained from multiple suppliers including ENZO Life Sciences (Farmingdale, NY, USA), ProSpec Bio, cat # ENZ-769 (East Brunswick, NJ, USA), Antibodies-Online, cat # ABIN7400042 (Limerick, PA, USA), Elabscience cat # PKSH033450 (Houston, TX, USA), Abcam cat # ab280347, ab174022 (Cambridge, UK). MMP-2 enzyme inhibition kits were from ENZO Life Sciences (Farmingdale, NY, USA). The human U87MG glioblastoma cells were from American Type culture Collection (ATCC) cat # HTB-14 (Manassas, VA, USA). Recombinant Neuropilin–1 (NRP–1, UniProt: O14786) was obtained from ACROBiosystems, cat #NR1-H5228 (Newark, DE, USA), containing amino acid residues Phe22-Lys644, with a polyhistidine tag at the C-terminus. For cell culture, DMEM high glucose cat # 11965092, Trypsin EDTA (1X) cat # 25200-072, PBS pH 7.4 without Ca2+ or Mg2+ ions cat # 10010-023, from Life Technologies (Carlsbad, CA, USA).

### Computational Methods

#### Structure Preparation

The structure of Ctx for docking was the first conformation of the NMR solution structure from the Protein Data Bank (PDB 1CHL) [45,46]. MMP-2 structure for docking was the lowest energy structure from the free energy landscape in Voit-Ostricki *et al.* 2019 [10]. PDB structures of four C-terminal fragment peptides (Table 2) was obtained by truncating the structure of Ctx, and in opposite to fragmentation to Dastpeyman and colleagues [44]. we introduced isosteric Ser substitutions for those Cys residues that are not in disulfide bridges. Cys substitutions were introduced to form non-native disulfide bridges. *N*- and *C*-terminal residues were acetyl and amide protected, respectively.

**Table 2.** Designed peptide fragments of Ctx.

| Peptide analogs | Amino acid sequence* |
| --- | --- |
| P75 | Ac-[Ser<sup>35</sup>,Cyc(28,33)]Ctx(24-36)-NH<sub>2</sub> |
| P76 | Ac-[Cys<sup>26</sup>,Ser<sup>28,33</sup>,Cyc(26,35)]Ctx(24-36)-NH<sub>2</sub> |
| P77 | Ac-[Cys<sup>24</sup>,Ser<sup>28,33</sup>,Cyc(24,35)]Ctx(24-36)-NH<sub>2</sub> |
| P78 | Ac-[Cyc(28,33)]Ctx(27-34)-NH<sub>2</sub> |

\*Ac and -NH<sub>2</sub>, are acetyl and amide, protecting groups respectively.

#### Molecular Docking and Molecular Dynamics Simulation

PDB structure of Ctx and its peptide fragments together with MMP-2 were submitted to molecular docking simulations using HPEPDOCK [47], HADDOCK [48], and AlphaFold2 (AF2) [49] methods.

##### Molecular Docking

For the HPEPDOCK method, PDB structure of MMP-2 and primary structure of peptides were submitted to the web server [50] using the default parameters. From the 100 poses, the four poses with the best docking scores were accepted for the Ctx – MMP-2 complex, and ten were accepted for each of the peptide – MMP-2 complexes.

Using the HADDOCK targeted docking method, the PDB structure of both MMP-2 and each peptide were submitted to the web server [51]. This docking methodology yielded ten clusters each containing four poses. The pose from the cluster with the smallest weighted sum of the HADDOCK scoring function was accepted for each of the peptide – MMP-2 complexes.

AF2 [52] dockings were executed with a local copy of ColabFold [52] using the primary structures of MMP-2 and the peptides as inputs. Five optimized poses were generated for each of the peptide – MMP-2 complexes and the pose with the highest rank was used for further structural analysis. Predicted template modeling scores and interface predicted template modeling scores were used to assess the quality of overall structure prediction and docking [49].

##### Molecular Dynamics Simulations

The structure of accepted poses from each docking were prepared for simulations by adding missing hydrogens and terminal protections, as appropriate, using the YASARA program [53,54]. The complexes underwent energy minimization without and with water molecules using the AMBER-FB15 force field [55] and Particle Mesh Ewald electrostatics method [56] for periodic boundary conditions. The resultant structures were then submitted to molecular dynamics (MD) simulations using the GROMACS2022 software package [57] and the CHARMM36m force field [58]. Complexes were solvated with TIP3P water [59] in a dodecahedron box so that the distance between the edge of the box and the protein-peptide complex was 1.2 nm. Net charges were neutralized with Na+ and Cl- ions, and the final concentration of NaCl was set to 150 mM. [59]. After 1000 steps of steepest-descent energy minimization, 1 ns constant number of particle, volume, and temperature (NVT) MD simulation was performed at 300 K using the stochastic velocity rescale method [60] using temperature coupling constants (τT) of 0.1 ps. Positions of backbone atoms were restrained with 1000 kJ mol^-1^ nm^-1^ force constant and MD integration time-step was 0.002 ps. Subsequently, 10 ns equilibration was performed at 310 K and a pressure of 1 atm (NPT) using the stochastic velocity rescale method [60] and Berendsen barostat [61] respectively. The isothermal compressibility was 4.5⋅10^-5^ bar ^−1^ and the pressure coupling constant (τP) was 2 ps. Hydrogen atoms were constrained using the Lincs algorithm [62]. Next, a full production of a 100 ns NPT simulation was performed using the Parrinello-Rahman barostat [63] at 1 atm and 310 K. Time constants τT and τP were 1 ps and 5 ps, respectively. Neighbor searching used the Verlet cutoff scheme [64]. Particle-Mesh-Ewald electrostatics [56] used a cutoff distance of 1.2 nm, and Fourier spacing was 1.2 nm. Van der Waals interactions were cut off at 1.0 nm and 1.2 nm at the short and long range, respectively, with force-switch enabled.

Because multiple poses were accepted for each peptide – MMP-2 complex from the HPEPDOCK docking method, binding poses were rescored to select the lowest energy complexes for further MD simulations. The binding energies (ΔEb) of each peptide to MMP-2 were calculated in the last 50 ps of each trajectory using gmx_MMPBSA method [65,66]. The resultant lowest energy complexes represented the top complexes from the HPEPDOCK pool. The peptide – MMP-2 complexes with the most negative ΔEb from each docking method were submitted to a 500 ns MD simulation using the Parrinello-Rahman barostat and the same parameters as above.

#### Trajectory Analysis

For each peptide – MMP-2 complex, movement of Cα-atoms were calculated using covariance matrices generated by the GROMACS *covar* module. To determine if equilibration had been achieved two methods were used. (1) Using eigenvectors corresponding to the 150 highest eigenvalues and sampling the trajectories at 2 ns configurational entropy was calculated as a function of time [67–69]. (2) The overlap of the sampled region of subspace with a 2 ns sampling frequency was determined using the method of Hess [69]. The GROMACS *rms* module was used to calculate the root mean square deviation (RMSD) of Cα-atoms for the 500 ns simulation trajectory. Trajectories were submitted to clustering using the GROMOS method [70]. The gmx_MMPBSA tool was used to calculate binding energies (ΔEb) and residue decomposition of peptides to MMP-2 during the last 50 ps of the MD trajectories. Decomposition of residues was considered for distances fewer than 6 Å between the atoms of the interacting residues.

### Experimental Methods

#### Wound-Healing Assay

The U87MG GBM cell line (U-87 MG, ATCC® HTB-14) was cultured in Dulbecco’s Modified Eagle Medium (DMEM) and appropriate growth factors at 37°C in a humidified 5% CO2 atmosphere. Cells were seeded at 40,000 cells per well in 96-well plates and allowed to adhere overnight. Then, mitomycin C (3 μg/ml) was added to inhibit further cell replication, and 50 µM Ctx and its analogs were added. Untreated wells served as a control. Wounds were created by manually scratching the cell monolayer with a 200 μL multi-channel pipette tip, and plates were analyzed at 0 h and 24 h time points using the MetaXpress image acquisition software (Molecular Devices, San Jose, CA). The ImageJ Ver. 1.54f program (NIH (National Institutes of Health) was used for image analysis. Wound closure was quantified using the *Wound_Healing_Size_tool_plugin.*, analyzing the percentage of wound area covered by migrating cells. To determine significance, statistical analysis was performed using appropriate methods (t-test or ANOVA) using GraphPad Prism software, version 10.0.3.

#### MMP-2 Inhibition Assay

The MMP-2 inhibition assay was performed using the Enzo Life Sciences MMP-2 drug discovery kit. and the quenched fluorogenic substrate OmniMMP RED. Reference substrate was diluted to a final concentration of 5 μM, while the MMP-2 enzyme was diluted to 58 mU/µL, and the N-Isobutyl-N-(4-methoxyphenylsulfonyl)glycyl hydroxamic acid (NNGH) inhibitor was diluted to a concentration of 1/200 in assay buffer. Solution of Ctx and analog peptides were prepared at concentrations of 100 μM, 50 μM, and 1 μM in water. The assay was performed using a black 96-well microplate. Fluorescence was measured at excitation/emission wavelengths of 545/576 nm, with a cutoff at 570 nm using Gen5 microplate reader (BioTeK, Shoreline, WA). Baseline fluorescence was recorded before adding NNGH inhibitor which was used as a positive control for MMP-2 inhibition. Reaction mixtures in the plate were incubated at 37 °C for 10 min, and fluorescence was measured at 10 min intervals for the first 30 min, then at 30 min intervals until 2 h had elapsed. Percentage of inhibition of MMP-2 activity by Ctx and its analogs was calculated as percent of control. Statistical analysis was performed using ANOVA with GraphPad Prism software, version 10.0.3.

#### Differential Scanning Fluorimetry (DSF)

The melting temperatures (Tm) of recombinant MMP-2 and its complexes with peptides were ascertained using BioRad CFX384 Touch real-time PCR (Hercules, CA, USA). The experiments were performed as previously described [71,72]. The unfolding of the protein was detected using the fluorescent SYPRO-orange dye (Invitrogen, Carlsbad, CA, USA). MMP-2 and prospective ligand peptides were dissolved in 7.1 pH phosphate buffered saline (154 mM NaCl, 5.6 mM Na2PO4, 1.1mM KH2PO4), and the reaction mixture was made to a final concentration of 1 µM of protein, either 30 µM (Ctx) or 50 µM (P75 and P76) of peptide, and 1000x dilution of the SYPRO-Orange dye stock from the manufacturer. Excitation and emission filters of the dye were set at 450−490 nm and 560−580 nm, respectively. The temperature for the reaction was held at 25 °C for 5 min, and then a gradient of 25 °C to 95 °C was applied at a rate of 0.3 °C/min, and the fluorescence was recorded. To determine Tm of the protein, the first derivatives of melting curves were plotted using GraphPad Prism software, version 10.0.3.

#### Surface Plasmon Resonance (SPR)

Surface plasmon resonance (SPR) reagents, plates, and the NTA chip were purchased from GE Healthcare Life Sciences, now Cytiva (Marlborough, MA, USA). Measurements were performed using the Biacore 8000 SPR System (Marlborough, MA. USA). The running buffer used for protein immobilization and peptide binding consisted of 20 mM Tris, 150 mM NaCl, 50 mM EDTA and 0.0005% Tween-20, adjusted to pH 7.4. MMP-2 and NRP-1 were immobilized onto the NTA chip at 15 nM and 25 nM concentration, respectively. Peptides were injected over the immobilized proteins using two-fold serial dilutions from 500 µM to 3.90 µM. Three experiments with four replicates of each were performed. The SPR response curves generated from the Biacore Insight Evaluation Software were fitted to a steady-state affinity model to obtain an equilibrium dissociation constant (KD) for each peptide — protein interaction.

## Results

### Computational Results

#### HPEPDOCK

Using 100 ns MD simulation and subsequent MM-PBSA calculations the rank order of docking poses for each peptides changed as follows: Ctx – MMP-2, pose 4; P75 – MMP-2, pose 8; P76 – MMP-2, pose 5; P77 – MMP-2, pose 10; P78 – MMP-2, pose 4 (Table 3). Configurational entropy calculated from 500 ns simulations trajectories of each of these selected poses sharply decreased during the first 50 ns and subsequently plateaued and the overlap of sampled region of subspace for each simulation after about 200 ns was above 0.5 indicating that the simulations reached equilibrium (S1 Fig). Cα RMSD analysis of these trajectories showed that structures of Ctx and P78 peptides had the least, while P76 and P77 had the most structural fluctuations in the complexes (S2 Fig). In the P75 – MMP-2, P76 – MMP-2, and P77 – MMP-2 complexes, the structure of MMP-2 hardly changed (S2 Fig). All complexes had a large negative ΔEb computed by gmx_MMPBSA method, indicating favorable interactions between Ctx and its analogs with MMP-2 (Table 4). Analysis of central structures of the largest clusters of each of the 500 ns trajectories (Fig 3) and residue-residue decomposition of ΔEb from MM-PBSA calculations (Fig 4) of each complex revealed that Ctx interacted with the collagenase-like 1 domain and hemopexin domain; P75 interacted with the collagenase-like 1 domain; P76 interacted with the collagenase-like 1 domain, collagen binding domain, and hemopexin domain; P77 interacted with the collagenase-like 1 domain; and P78 interacted with the collagenase-like 1 domain, collagenase-like 2 domain, and hemopexin domain.

**Fig 3.**
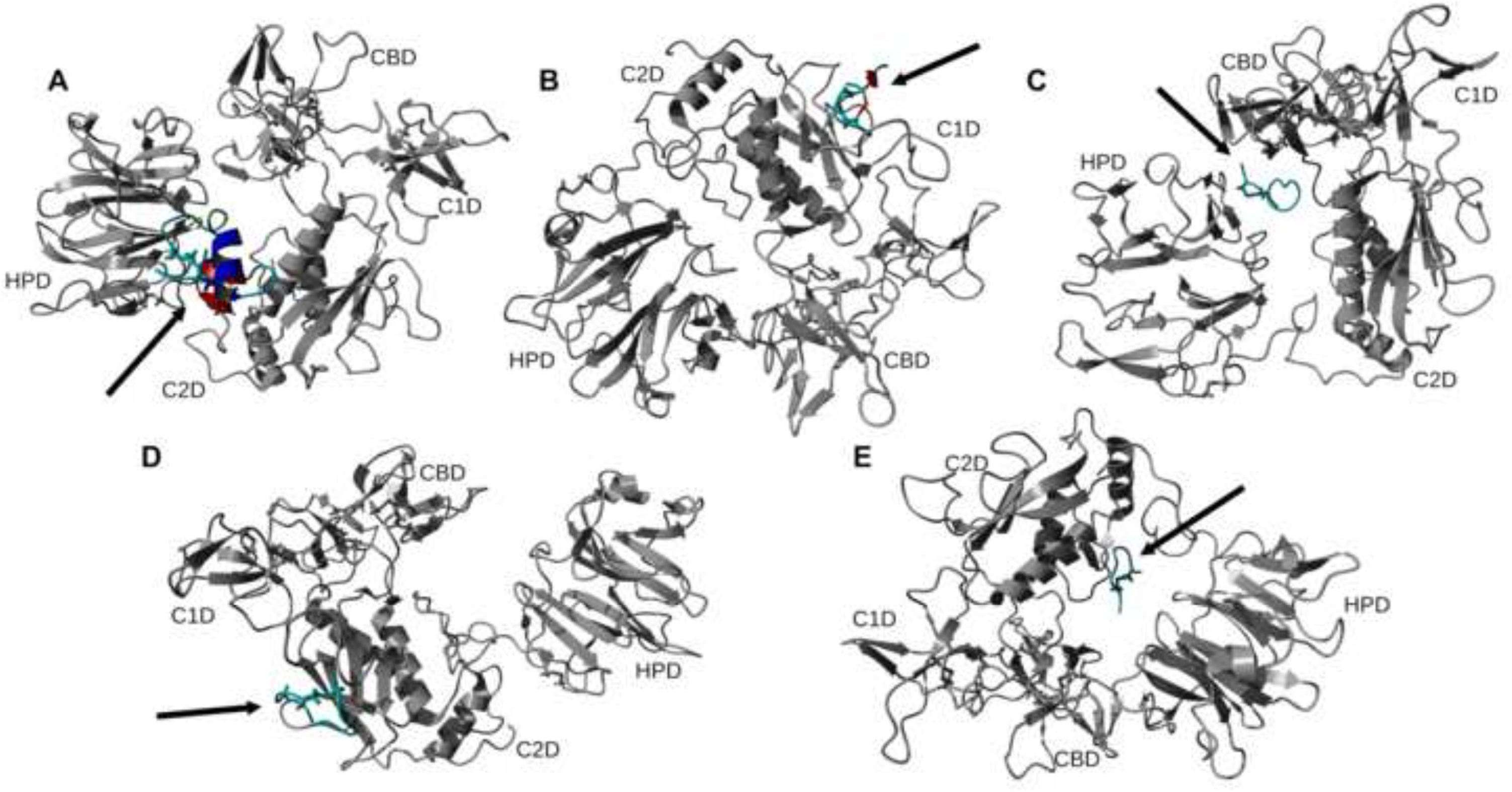
Ribbon representation of the central structures of the largest clusters of 500 ns trajectories of peptide – MMP-2 complexes using the re-ranked top HPEPDOCK docking poses. The orientation of MMP-2 has been optimized to allow for viewing of the peptide-MMP-2 interactions. Peptide ligand regions identified as follows: random coil, cyan; alpha helix, blue; beta sheet, red; beta turn, green. Disulfide bridges are shown in sticks. For MMP-2 receptor: collagenase-1 region, orange; collagenase-2 region, purple; remainder of protein, gray. Abbreviations for MMP-2 domains are from Table 1.

**Fig 4.**
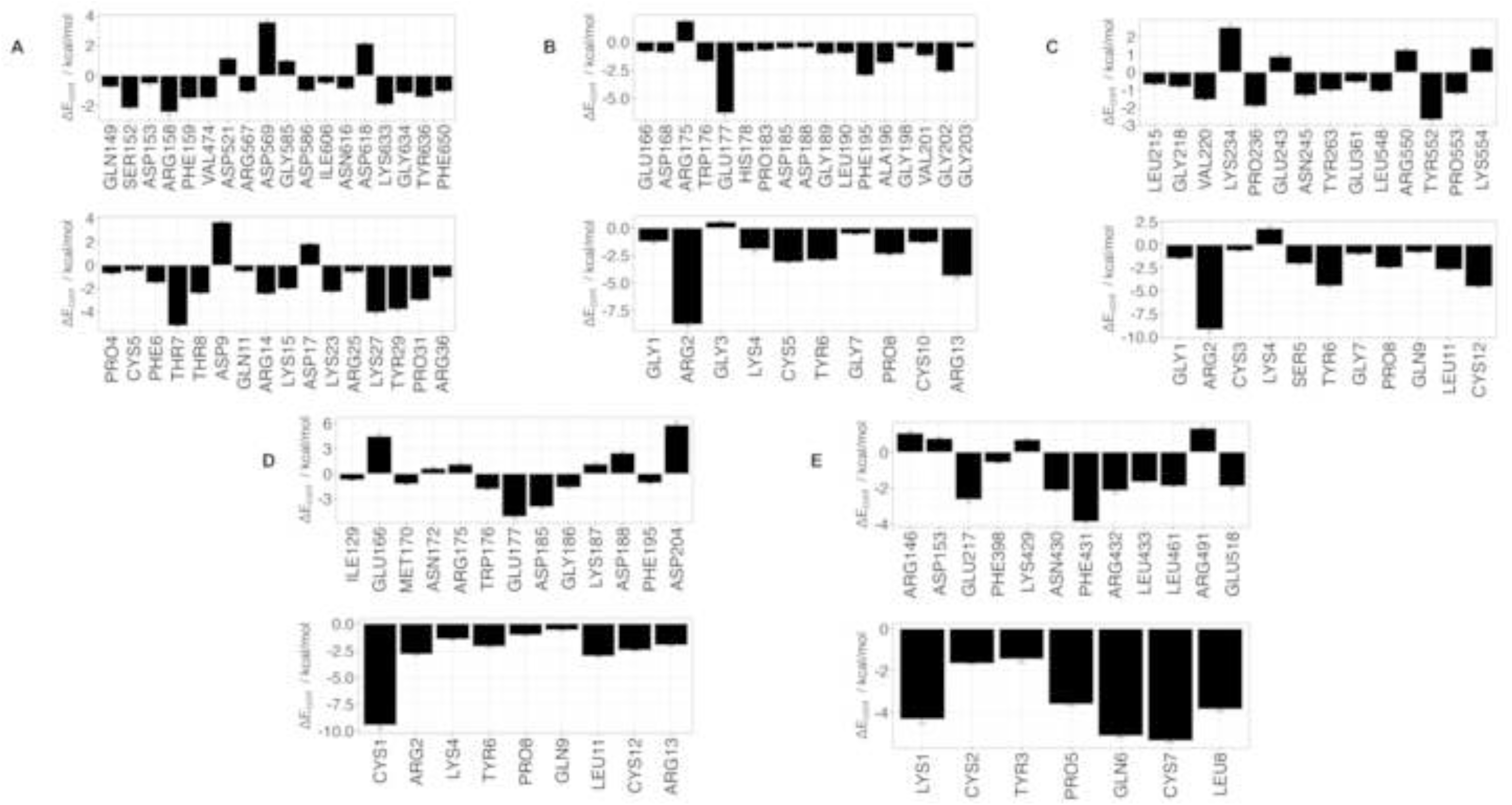
Residue contribution to ΔEb from MM-PBSA calculations for peptides and MMP-2 complexes obtained from the 500 ns trajectories MD simulations. (**A**) Ctx – MMP-2; (**B**) P75 – MMP-2; (**C**) P76 – MMP-2; (**D**) P77 – MMP-2; (**E**) P78 – MMP-2. Top panel contains residues from MMP-2, bottom panel contains residues from peptides. ΔEcont is the contribution to binding energy for each residue.

**Table 3.**
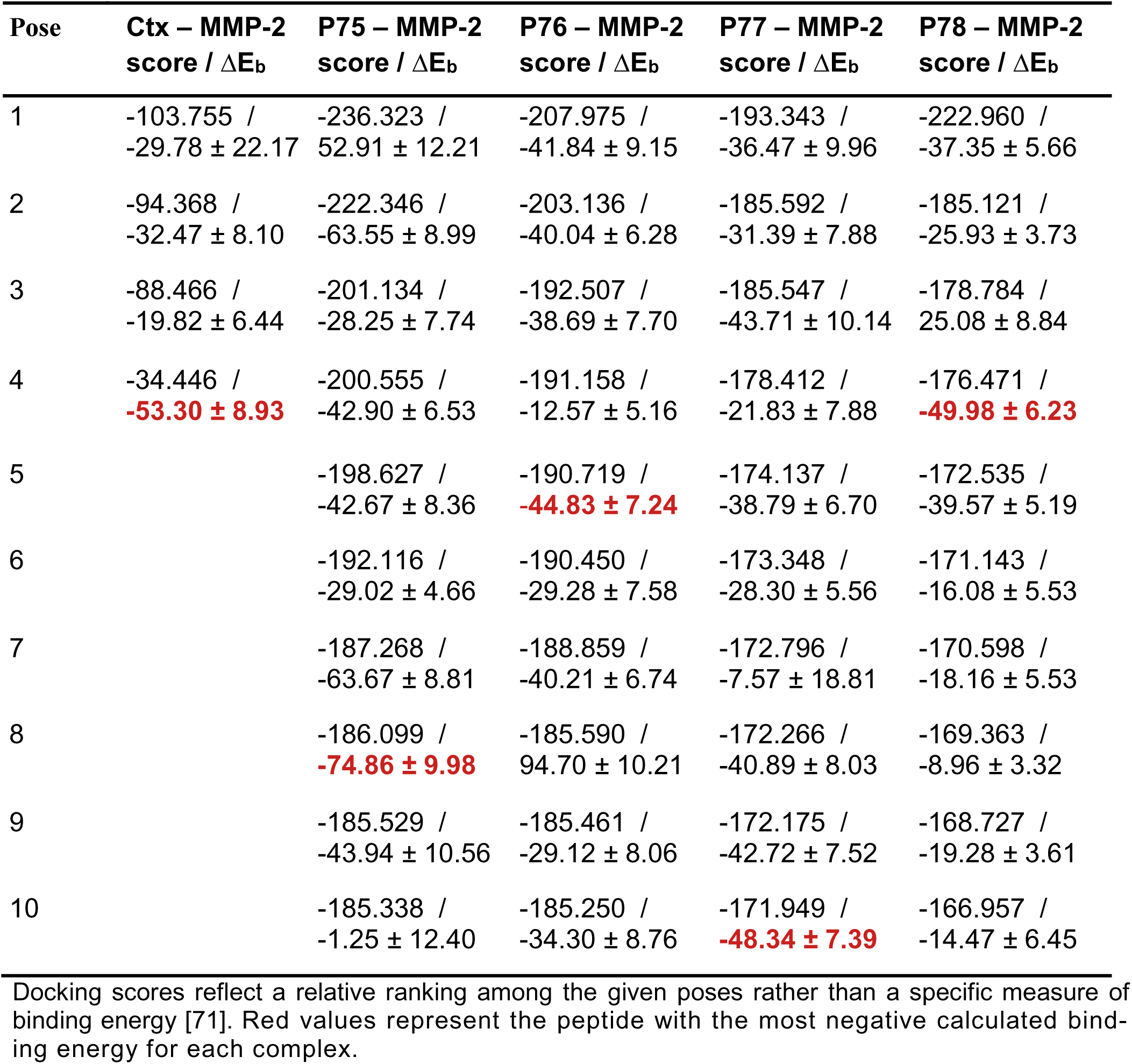
Docking scores and binding energy (ΔEb, kcal/mol) after 100 ns molecular docking of accepted HPEPDOCK poses.

**Table 4.** Binding energy (ΔEb) of peptides among docking methods.*

| Peptide | HPEPDOCK<br>(kcal/mol) | HADDOCK<br>(kcal/mol) | AlphaFold2<br>(kcal/mol) |
| --- | --- | --- | --- |
| Ctx | **-62.53 ± 7.2** | -37.82 ± 7.9 | -53.02 ± 11.3 |
| P75 | **-61.65 ± 6.7** | -21.80 ± 6.9 | -58.49 ± 10.3 |
| P76 | -50.72 ± 9.6 | **-52.16 ± 7.3** | -51.85 ± 9.3 |
| P77 | -37.59 ± 10.0 | -27.78 ± 6.5 | **-72.89 ± 11.2** |
| P78 | **-51.72 ± 6.4** | -41.90 ± 5.4 | -29.94 ± 11.0 |

\*Red values represent the peptide with the most negative calculated binding energy for each docking method.

#### HADDOCK

Docking scores for peptide − MMP-2 complexes obtained with the HADDOCK methodology are listed in Table 5. The docking score in this method is a weighted sum of the calculated complex energies. The top structures from the clusters with the smallest docking scores for each of the peptide – MMP-2 complexes were selected for MD simulation. Configurational entropy calculated from 500 ns simulation trajectories of each of these selected poses sharply decreased during the first 50 ns and subsequently plateaued. Except for the P76 − MMP-2 complex simulation (S3C Fig), the overlap of sampled region of subspace for the other simulations continuously increased and after about 200 ns was above 0.5 indicating that the simulations reached equilibrium. For the P76 – MMP-2 complex simulation, between 200 ns to 400 ns the overlap of sampled region of subspace remained constant ∼ 0.4, but after that steadily increased indicating that the simulation reached the equilibrium (S3 Fig). Cα RMSD analysis of the trajectories showed that the structure of Ctx in the complex was the most stable (S4 Fig). The structure of P75 fluctuated most between ∼ 0.5 nm and 5 nm indicating a loose binding. After 350 ns, the structure of P77 together with MMP-2 experienced a major structural transition before reaching an equilibrium (compare to the overlap of sampled region of subspace in S3C Fig). In the MMP-2 − P75 and − P78 complexes the structure of MMP-2 hardly changed, while the structure of P75 fluctuated from 0.4 nm – 5 nm indicating that the peptide at this pose did not form stable complex. All complexes had negative ΔEb computed by gmx_MMPBSA utility, indicating favorable interactions between the peptides and MMP-2 (Table 4). Analysis of representative structures of the largest clusters of each of the 500 ns trajectories (Fig 5) and residue-residue decomposition of ΔEb from MM-PBSA calculations of each complex (Fig 6) revealed that only P76 interacted within 6 Å of the residues of MMP-2. The most favorable interaction was with the collagen binding domain. Although not interacting with MMP-2, Ctx was positioned closest to the collagenase-like 2 domain, P75 was positioned closest to the collagenase-like 2 domain, P77 was positioned closest to the collagen binding domain, and P78 was positioned closest to the collagenase-like 1 domain.

**Fig 5.**
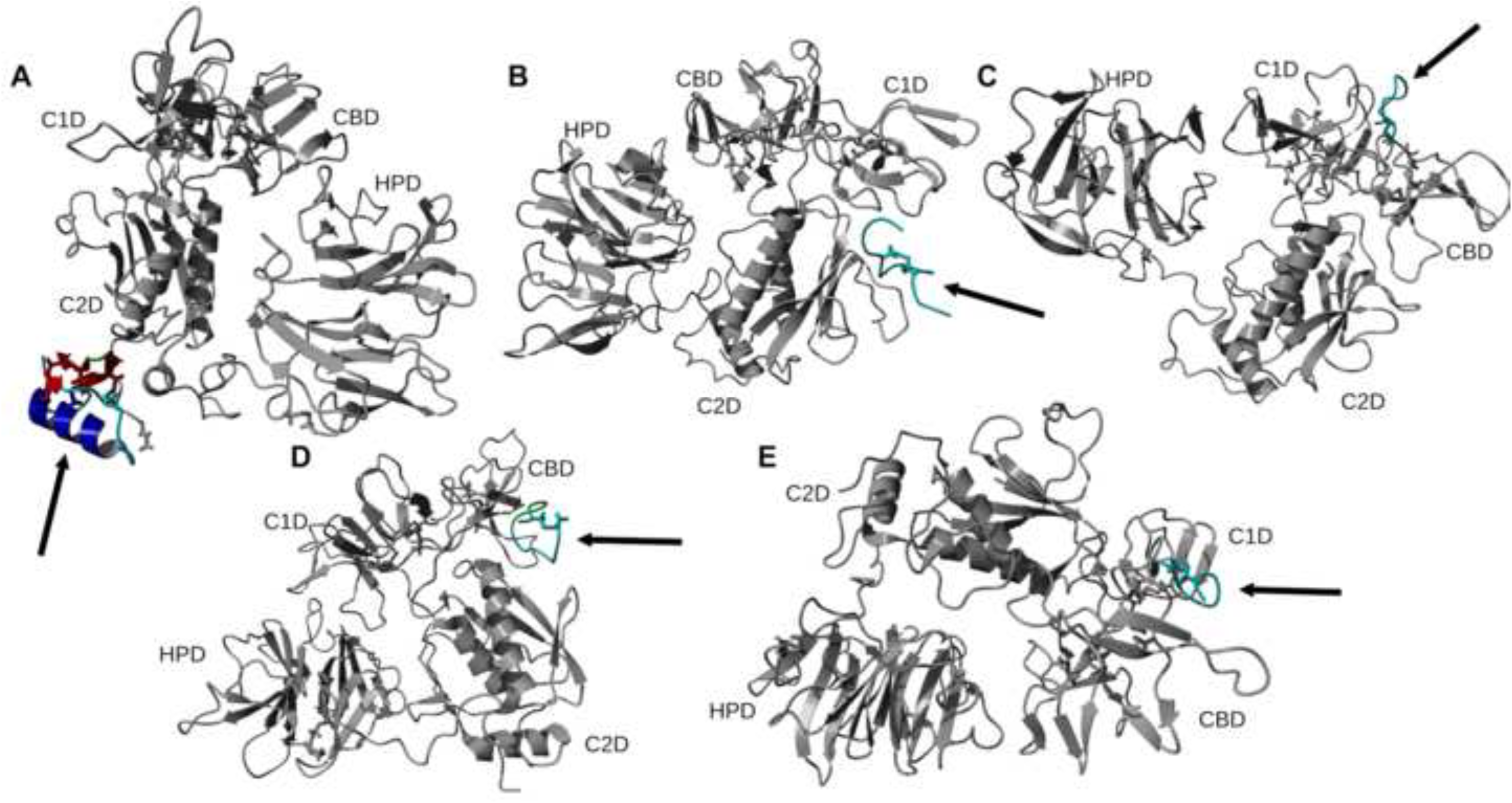
Representative structures of the largest clusters of 500 ns trajectories of peptide – MMP-2 complexes obtained with the HADDOCK method. The orientation of MMP-2 has been optimized to allow for viewing of the peptide-MMP-2 interactions. Protein ligand regions identified as follows: random coil, cyan; alpha helix, blue; beta sheet, red; beta turn, green. Disulfide bridges are shown in sticks. For MMP-2 receptor: collagenase-1 region, orange; collagenase-2 region, purple; remainder of protein, gray. Abbreviation for MMP-2 domains are from Table 1.

**Fig 6.**
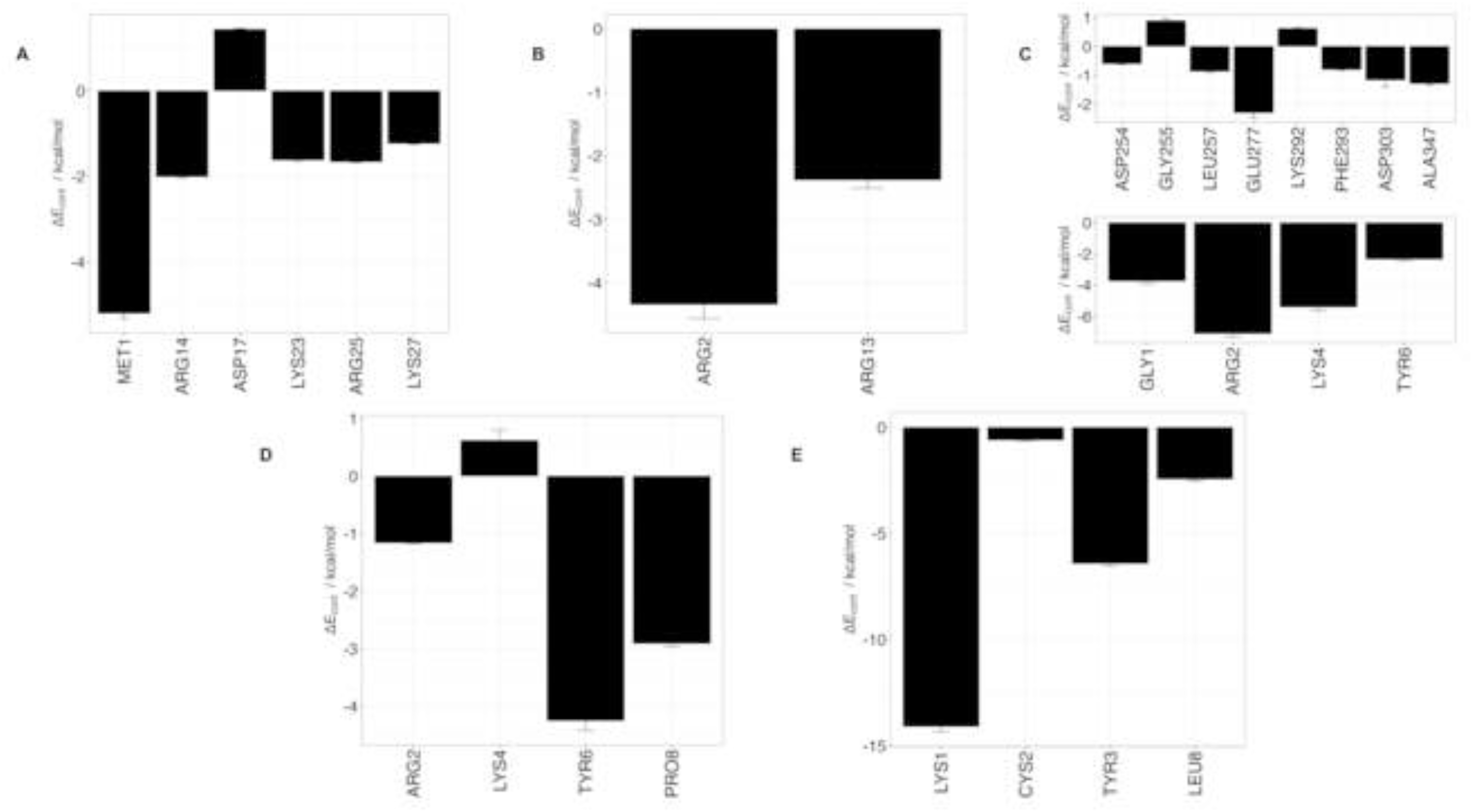
Residue-residue decomposition of ΔEb from MMP-BSA calculations of peptides and MMP-2 complexes obtained with the HADDOCK method. (A) Ctx – MMP-2; (B) P75 – MMP-2; (C) P76 – MMP-2; (D) P77 – MMP-2; (E) P78 – MMP-2. Top values contain residues from MMP-2, bottom values contain residues from peptides. For Ctx, P75, P77, and P78, there were no contributions from MMP-2 residues. ΔEcont is the contribution to binding energy for each residue.

**Table 5.** Docking scores of accepted HADDOCK poses.

| Peptide | HADDOCK score* |
| --- | --- |
| Ctx – MMP-2 | 117.5 ± 5.0 |
| P75 – MMP-2 | 127.0 ± 9.0 |
| P76 – MMP-2 | 135.6 ± 8.6 |
| P77 – MMP-2 | 148.8 ± -2.6 |
| P78 – MMP-2 | 180.6 ± 5.1 |

\*Smallest weighted sum of the HADDOCK scoring function

#### AF2

Modeling scores for each of the poses for the peptide − MMP-2 complexes obtained with the AF2 docking method are listed in Table 6. All the selected poses had pTM scores greater than 0.8 and ipTM scores greater than 0.68 therefore, these poses most likely represent accurate binding predictions. Configurational entropy calculated from 500 ns simulation trajectories of each of these selected poses sharply decreased during the first 50 ns and thereafter stayed nearly constant (S5 Fig). The subspace overlap for each simulation after about 200 ns was above 0.5, again indicating that the simulations reached equilibrium (S5 Fig). Cα RMSD analysis of each trajectory showed that peptide conformations remained relatively stable, but the conformation of MMP-2 fluctuated significantly (S6 Fig). Similar structural fluctuation was observe in ref 10. Each peptide − MMP-2 complex had a negative ΔEb that was comparable to that of the other two docking methodologies (Table 4). Analysis of representative structures of the largest clusters of each of the 500 ns trajectories (Fig 7) and residue-residue interaction decomposition of ΔEb from MM-PBSA calculations of each complex (Fig 8) revealed that Ctx interacted with the collagenase-like 1 domain, collagen binding domain, and collagenase-like 2 domain; P75 interacted with the collagen binding domain and hemopexin domain; P76 interacted with the collagen binding domain and hemopexin domain; P77 interacted with the collagen binding domain and hemopexin domain; and P78 interacted with collagenase-like 1 domain, collagen binding domain, and collagenase-like 2 domain.

**Fig 7.**
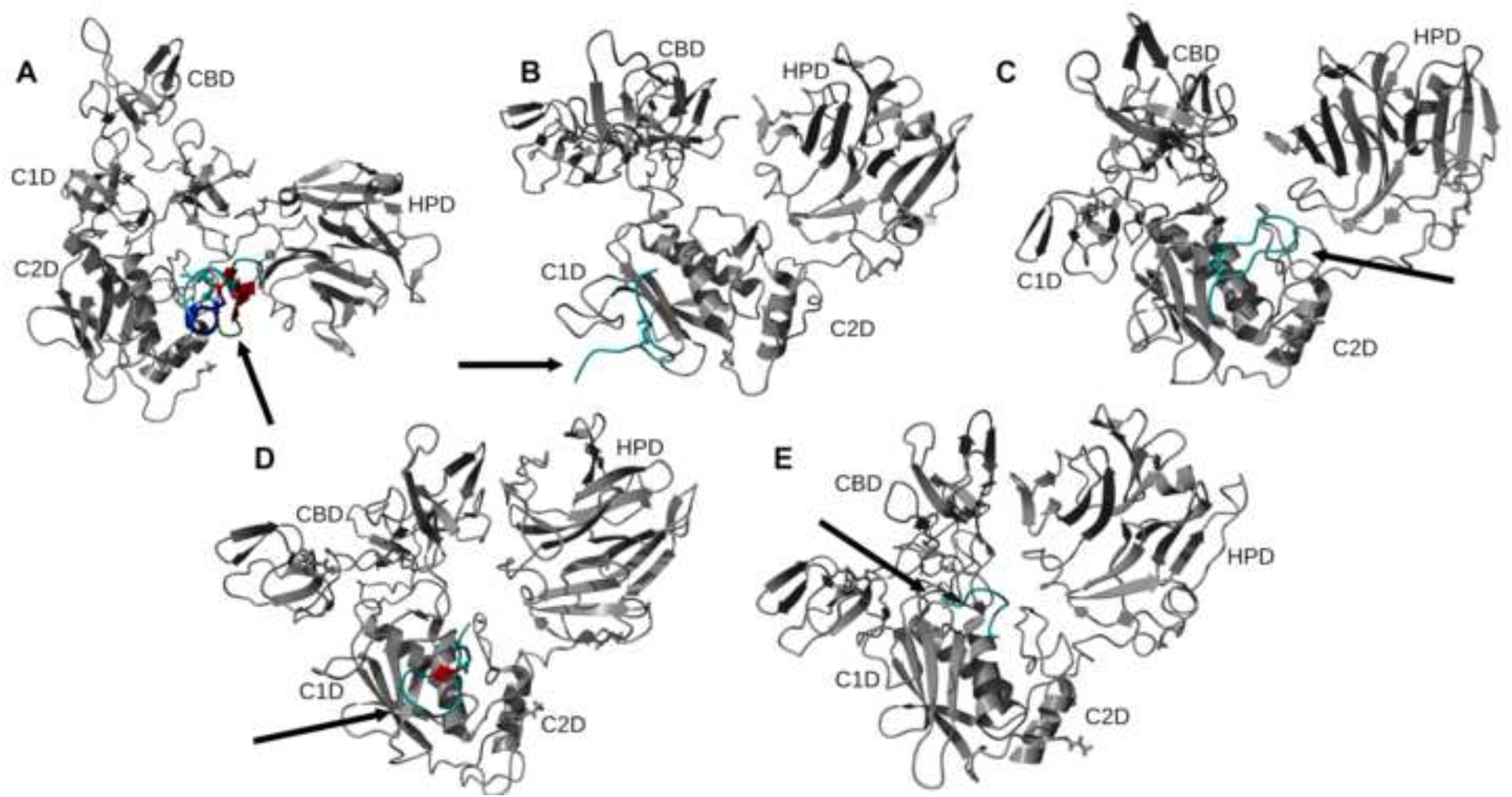
Representative structures of the largest clusters of 500 ns trajectories of peptide – MMP-2 complexes obtained with the AF2 method. Protein ligand regions identified as follows: random coil, cyan; alpha helix, blue; beta sheet, red; beta turn, green. Disulfide bridges are shown in sticks. For MMP-2 receptor: collagenase-1 region, orange; collagenase-2 region, purple; remainder of protein, gray. Abbreviation for MMP-2 domains are from Table 1.

**Fig 8.**
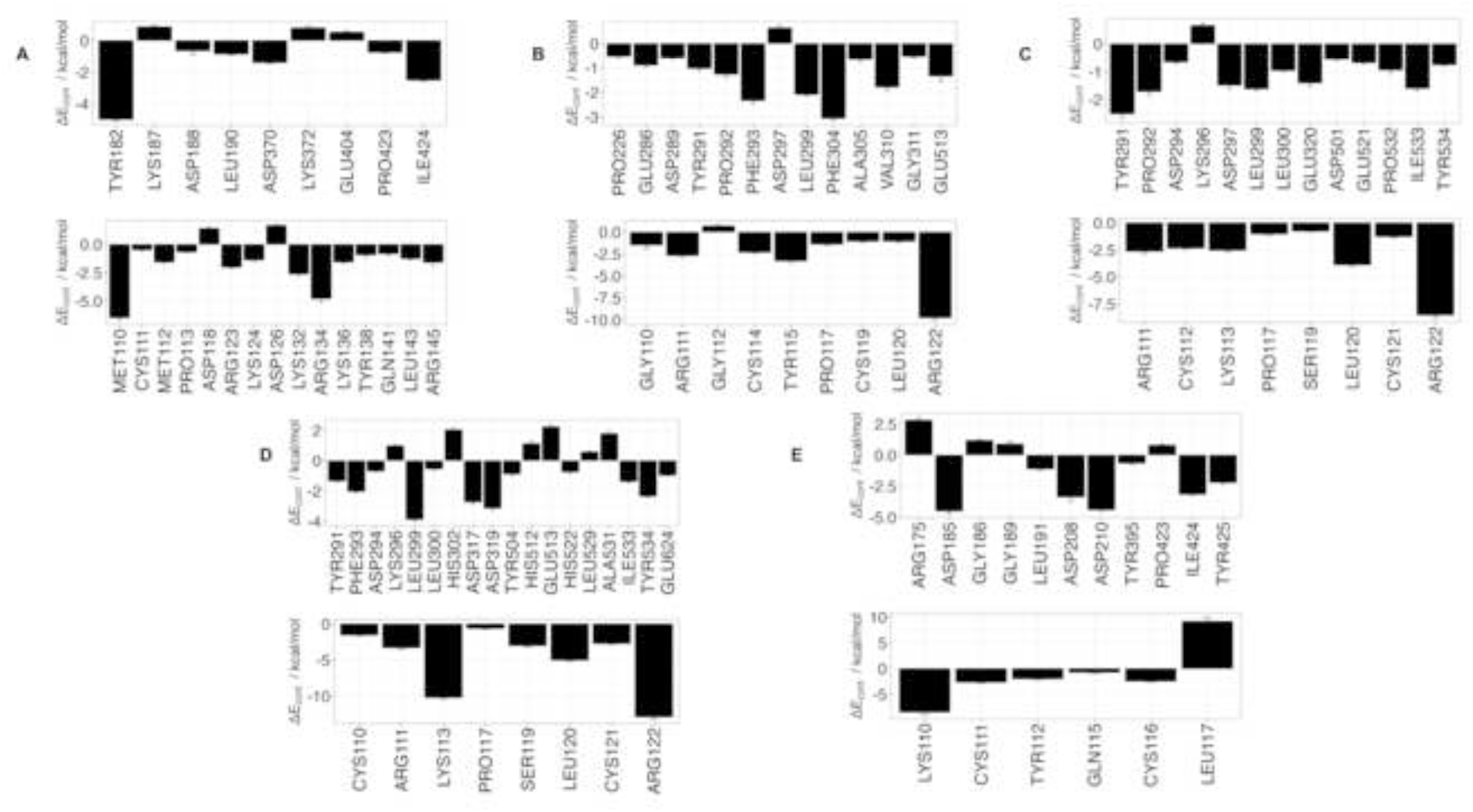
Residue-residue decomposition of ΔEb from MM-PBSA calculations of peptides − MMP-2 complexes obtained with the AF2 method. (A) Ctx – MMP-2; (B) P75 – MMP-2; (C) P76 – MMP-2; (D) P77 – MMP-2; (E) P78 – MMP-2. Top values contain residues from MMP-2, bottom values contain residues from peptides. ΔEcont is the contribution to binding energy for each residue.

**Table 6.** Confidence scores* of accepted poses obtained with the AF2 method.

| Complex | pTM | ipTM |
| --- | --- | --- |
| Ctx – MMP-2 | 0.823 | 0.617 |
| P75 – MMP-2 | 0.870 | 0.761 |
| P76 – MMP-2 | 0.870 | 0.770 |
| P77 – MMP-2 | 0.867 | 0.762 |
| P78 – MMP-2 | 0.829 | 0.621 |

\*pTM, predicted template modeling score; ipTM, interface predicted template modeling score

During MD simulation following all of the three docking methodologies, the residues from the receptor participating in binding were sparse and scattered. The collagenase-like 1 domain residues participating in binding are Asp153, Glu166, Arg175, Glu177, Trp176, Asp185, Lys187, Gly186, Asp188, Gly189, Leu190, and Phe195. Residues Tyr291, Pro292, Phe293, Asp294, Lys296, Asp297, Leu299, and Leu300 correspond to the collagen binding domain. Hemopexin domain residues are Pro423, Ile424, Glu513, Ile533, and Tyr534.

### Experimental Results

#### DSF

DSF was used to determine binding of the peptide to MMP-2. Thermal unfolding profiles of MMP-2 and peptide – MMP-2 complexes, and the corresponding first derivative curves are shown in Fig 9. The Ctx peptide did not cause a shift in melting temperature (ΔTm) of the protein (Table 7). P75 and P76 peptide led to ΔTm of 0.76 ± 1.56 and 0.63 ± 0.36, respectively. The lack of either a shift in Tm or the small change in Tm with large comparable SD indicates that Ctx and its analog did not interact with MMP-2. Furthermore, the results suggest the possibility that in studies that reported Ctx and its analogs binding to MMP-2, Ctx could be interacting with an entirely different target protein than MMP-2.

**Fig 9.**
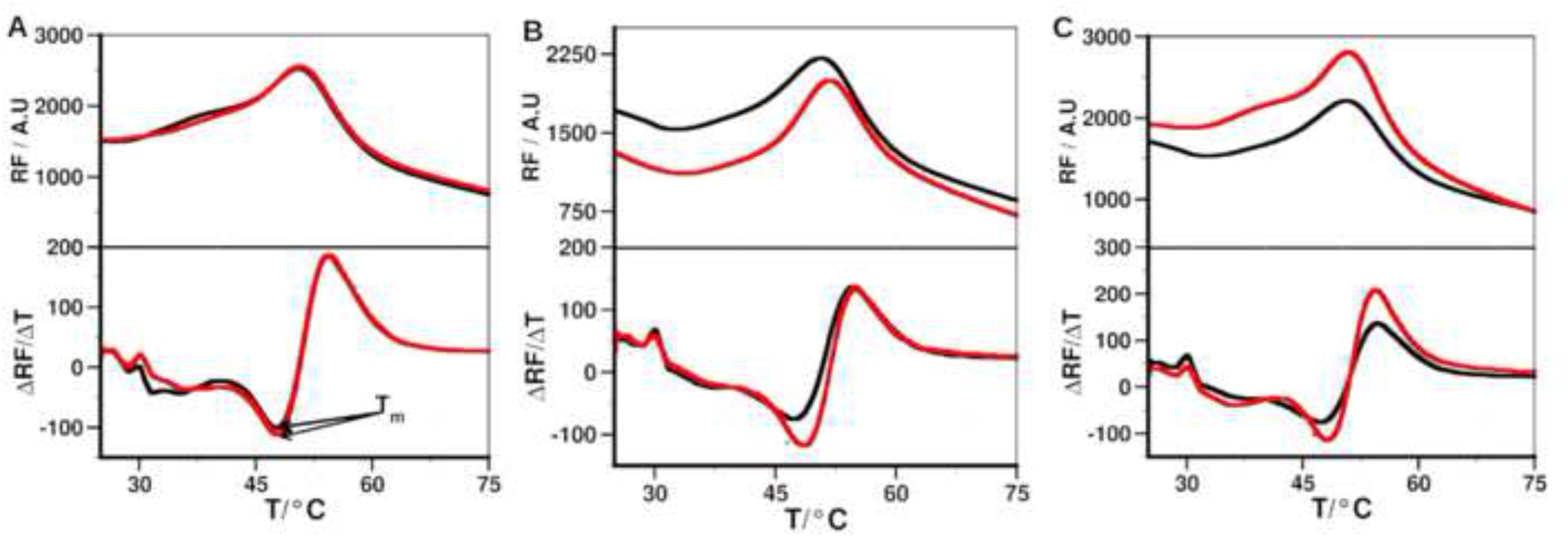
Binding of the peptides to MMP-2 determined by DSF. Top panels show the melting curve of the MMP-2 without (black) and with peptide (red). Bottom panels show the first derivative of the melting curve, the minimum of which represents the melting temperature. (A) Ctx; (B) P75; (C) P76.

**Table 7.** Change in melting temperature (ΔTm*) due to peptides binding to MMP-2.

| Peptide | $\Delta T_m \pm SD / ^\circ C$ |
| --- | --- |
| Ctx | $0 \pm 0$ |
| P75 | $0.76 \pm 1.56$ |
| P76 | $0.63 \pm 0.36$ |

\* $\Delta T_m$ values correspond to the average and SD of $n \geq 3$ .

#### SPR

Ctx and its fragments did not bind to MMP-2. In contrast, the binding affinity of the peptides for NRP-1, an alternative potential Ctx target [35], was measured. Both P78 and P75 weakly bound in the high micromolar range, with dissociation constants (KD) of 267 ± 103.02 µM and 330 ± 145.66 µM, respectively (S7 Fig). These results support the hypothesis that Ctx and its fragments do not possess a specific binding site on MMP-2 and may interact with an alternative target such as NRP-1.

#### MMP-2 Enzyme Inhibition Assay

MMP-2 Drug Discovery Kit with a quenched fluorogenic substrate, OMNIMMP, was used. Full-length Ctx was tested at concentrations of 1 µM, 5 µM, 10 µM, 15 µM, and 20 µM. Our results indicate that Ctx, at all tested concentrations, did not inhibit MMP-2 activity compared to the control inhibitor (Fig 10A). Similarly, Ctx fragments tested at 1 µM, 50 µM, and 100 µM also showed no inhibitory effect on MMP-2 activity (Fig 10B). The apparent lack of enzymatic inhibition by Ctx and its fragments suggests that they do not bind to the catalytic domain of MMP-2.

**Fig. 10.**
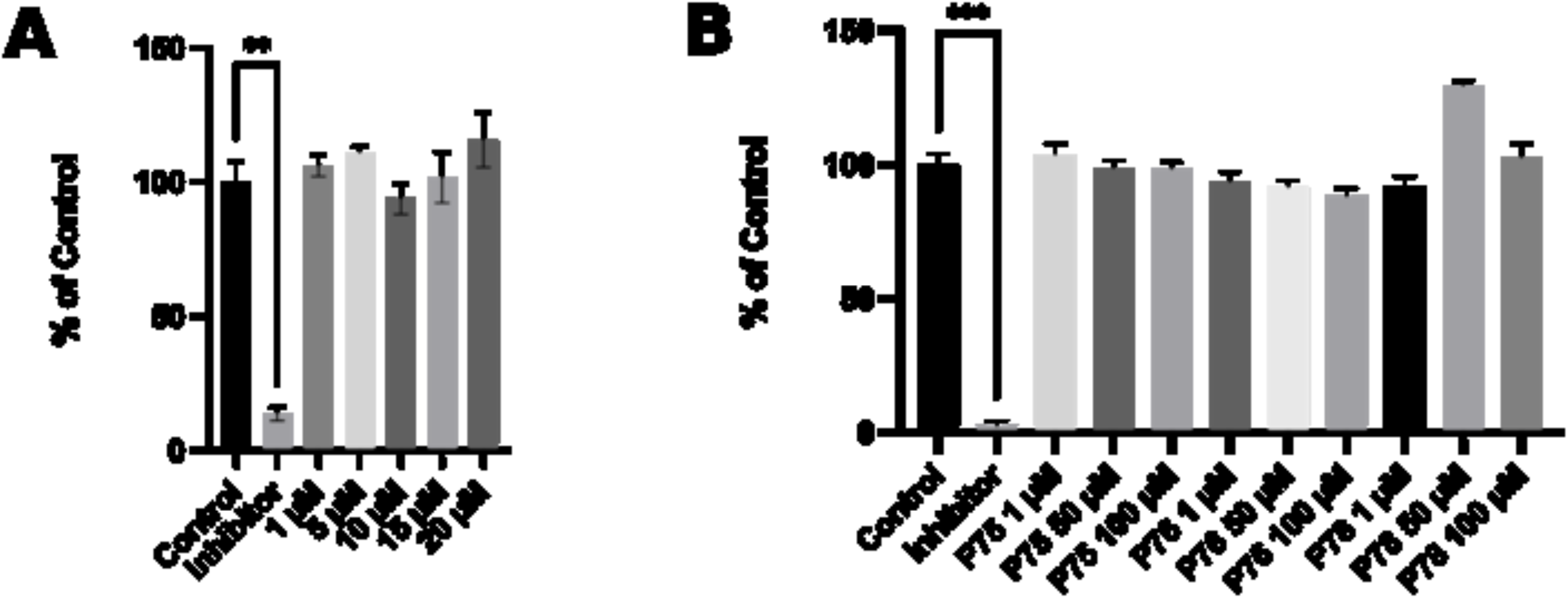
Inhibition of enzymatic activity of MMP-2. (A) Effects of Ctx at various concentrations on MMP-2 enzymatic activity. Data is presented by mean ± SEM and is not statistically significant. (B) Effects of Ctx, P75, P76, and P78 on MMP-2 enzymatic activity, along with the NNGH positive control. Data is presented by mean and SEM. Compared to the control, the experimental groups are not statistically significant (one-way ANOVA).

#### Wound-healing Assay

The wound-healing assay was performed to examine the effect of Ctx, P75, P76 and P78 peptides on U-87MG cell migration. Ctx, P75 and P76 showed no or little inhibitory effect on U-87MG cell migration compared to the control inhibitor in Fig 11. The inhibitory effects on U-87MG cells by the peptides is also visualized in S8 Fig. In contrast, P78 significantly inhibited cell migration of U-87MG cell by about 30%, surpassing the efficacy of Ctx.

**Fig 11.**
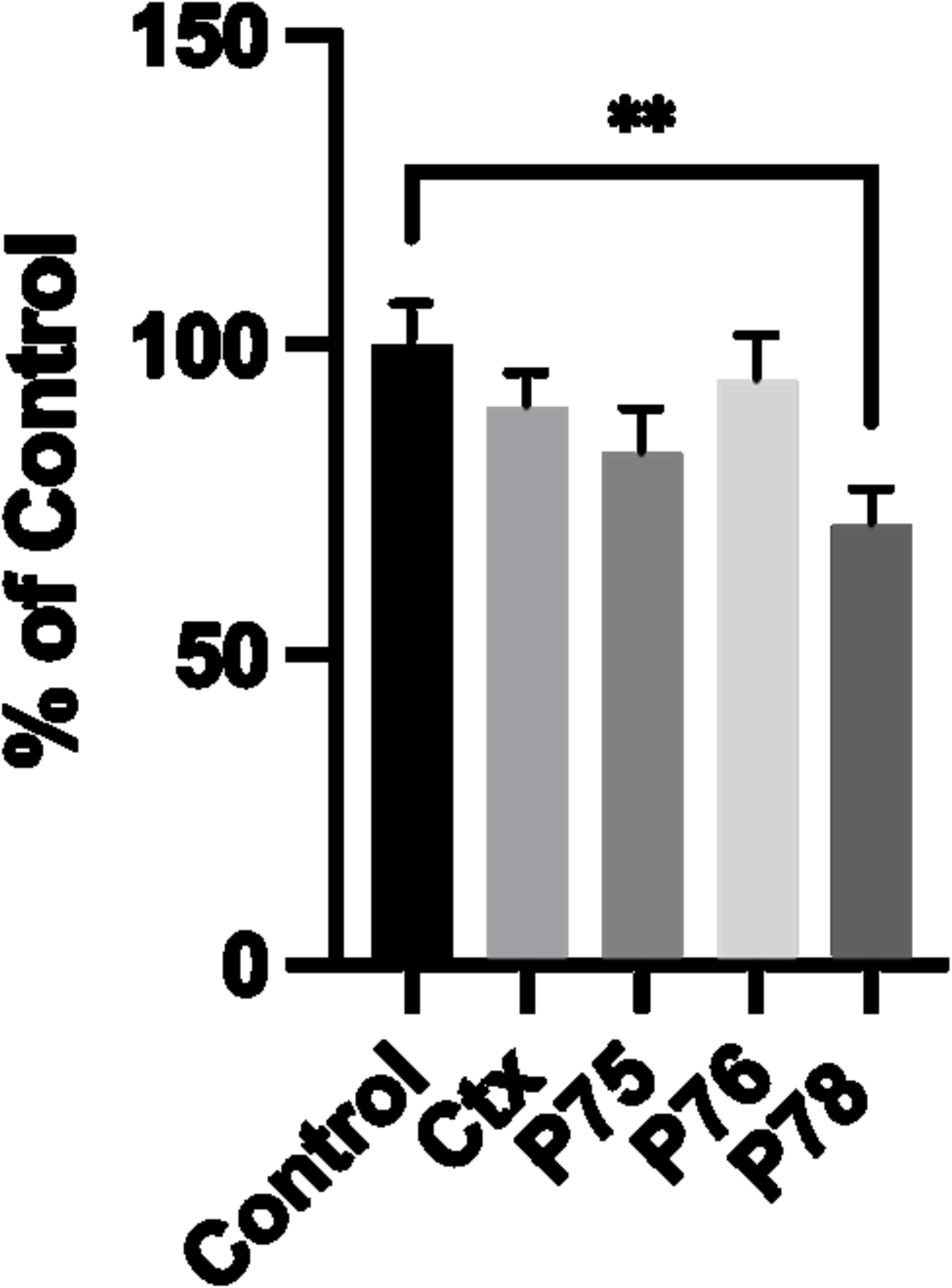
Wound-Healing Assay-U-87 MG cells. Quantification of cell migration inhibition using the wound-healing assay. Percentage of wound closure was measured using the Wound_Healing_Size_tool_plugin in ImageJ. Results are represented by the mean ± SEM. *\*\*\*P<0.0001 (One-way Anova)*

## Discussion

In the present work we used molecular docking and subsequent MD simulations to study Ctx and its peptide analogs binding to MMP-2. *In silico* methods were complemented by experimental (DSF and SPR) assays. Inhibition of MMP-2 enzymatic activity and inhibition of its glioblastoma cell migration by peptides were also assessed through an MMP-2 inhibition assay and a wound-healing scratch assay. Among the MD simulations of docking poses generated from HPEPDOCK had the most favorable binding energies except for P76 and P77, while HADDOCK had the least favorable binding energies except for P76 and P78. Decomposition of the peptide – MMP-2 complexes from each docking method reported overlap of participating residues of P75 – MMP-2 from HPEPDOCK and AF2, sharing agreement on Glu177, Pro183, Asp188, Leu190, Phe195, Ala196, Val201, and Gly202 from MMP-2. Complexes of MMP-2 with other peptides did not show any such consensus on active residues, and four of the five complexes generated using HADDOCK at the collagen binding domain revealed zero active residues from MMP-2. The agreement among the two blind docking methodologies suggests that Ctx and its fragments do bind to MMP-2, while the failure of targeted docking to show interactions suggests Ctx and its fragments bind to locations other than the collagen binding domain. This interpretation is supported by the DSF investigation showing no change in melting temperature, indicating no binding. Furthermore, investigation using SPR did not show binding as well.

There are several MMP-2 residues that contributed to binding, and some residues interacted with more than one peptide. These non-localized results further suggest that Ctx and its fragments are weak, nonspecific binders to MMP–2. Diaz and Suárez computed important substrate anchorage points of residues in MMP–2 involved in binding at the catalytic site, which when compared with our analysis finds shared residues Gly189, Leu191, Pro423, and Tyr425 [73]. However, those are the only such residues that contributed to binding during these simulations.

Together with the nonspecific binding representative clusters and absence of change in melting temperature, the present result cannot confirm the notion that Ctx binds to the three fibronectin regions in the collagen binding domain [33]. A more recent study using flow cytometry has demonstrated that Ctx binds to MMP-2 with a KD of 0.5 to 0.6 mM but has no statistically significant effect on the enzymatic activity of the enzyme [35]. This corroborates findings that Ctx binds allosterically to MMP-2 but does not bind to the catalytic site. Among the peptide fragments, the strongest binder to MMP-2 was P75 using the HPEPDOCK method, P76 using the HADDOCK method, and P77 using the AF2 method. The lack of inhibition of MMP-2 enzymatic activity by all peptides contrasts the claim by Deshane *et al.* that MMP-2 is the primary receptor for Ctx on GBM cell surfaces [28]. Inhibition of U-87MG cell migration by Ctx and the P75 and P78 fragments is consistent, however, with the notion that the C-terminal region of Ctx is most critical for inhibition of GBM migration [44]. These two peptides inducing inhibition fits within the context that P78 is a shortened copy of P75, especially since P78 was the most effective peptide overall in inhibiting migration. Although none of th e peptides experimentally inhibited the activity of MMP-2, this finding is consistent with Ctx having an alternative protein target within GBM cells.

This study does not establish a definitive binding location for Ctx to MMP-2. The presented findings suggest an allosteric mechanism, but further computational and experimental investigations are warranted. Alanine scanning or X-ray crystallography of the Ctx – MMP-2 complex would be possible methodologies for further study. Further, this study does not explain Ctx-induced inhibition of U-87MG migration despite the lack of inhibition of MMP-2 activity. Our results strongly suggest that alternative binding targets of Ctx should be evaluated for binding and inhibition, such as previous studies with neuropilin-1 or annexin A2 [35,74].

Our study clears the controversy concerning the role of Ctx with regard to MMP-2 expression in GBM. The unreliable binding of Ctx and its fragments to MMP-2 both *in silico* and *in vitro*, combined with inhibited migration of GBM *in vivo*, lay to rest the notion that Ctx-induced inhibition of MMP-2 is the responsible mechanism for GBM migration inhibition. The success of P78 in our wound-healing assay show that the C-terminal regions of Ctx are candidates for enzyme inhibition in future studies. In a previous study by Dastpeyman and colleagues [44] it was showed that a fragment containing the last eight residues of Ctx inhibited the U-87MG cell migration. Their peptide fragment, however, lacked the disulfide bridge and was somewhat less act ive. These findings suggest that the full Ctx peptide may not be necessary for inhibiting cell migration, and a limited set of residues may contribute to this biological activity. P78 emerged as the most effective peptide in inhibiting cell migration, highlighting its potential as a promising candidate for further investigation in therapeutic interventions targeting GBM cell migration. Future strategies may include antibody-drug conjugates, bispecific T-cell engagers, and CAR-T cell therapy [75].

Both SPR and DSF measurements support the idea that Ctx and its C-terminal fragments do not bind to MMP-2 at the catalytic site, which is seen by the absence of a substantial melting temperature shift in DSF and a lack of measurable KD in SPR. These results align with previous findings of Farkas *et al.*, which reported an inability to inhibit MMP-2 enzymatic activity by Ctx [35]. While earlier reports have shown that Ctx inhibits MMP-2, the exact mechanism of binding remains unknown. If Ctx acts as a non-competitive inhibitor, our findings would be consistent with the hypothesis that Ctx binds to MMP-2 through an allosteric mechanism. Peptides were assessed for their binding to NRP-1, another recently identified potential target of Ctx. The high micromolar KD values for both P75 and P78, as well as the substantial variability in the standard deviation, suggests that there is a weak binding interaction between the peptides and NRP-1. Further analysis is needed to determine the biological relevance of these interactions, as well as the potential involvement of other targets.

## Conclusion

This study provides insight into the reason that Ctx has been a promising strategy to reduce the aggression of GBM. Although we do not endorse Ctx as an inhibitor of MMP-2, this study supports that the C-terminal region of Ctx is responsible for binding to MMP-2 at regions separate from the catalytic domain. Inhibition of U-87MG migration by Ctx and its derived fragments established a link for Ctx to alternative protein targets expressed by GBM cells. The greater inhibitory activity of the polypeptide fragments further contributes as building blocks for more optimized inhibitors of GBM migration.

## Acknowledgments

Mass spectral analysis of synthetic peptides were performed by the Mass Spectrometry Core facility as a component of the Auditory Vestibular Technology Core within the Translational Hearing Center at Creighton University, School of Medicine.

## Supporting Information

**S1 Fig. MD simulations of the peptide – MMP-2 complexes using the re-ranked top HPEPDOCK docking poses.** Configurational entropy (black) and overlap of sampled region of subspace (red) of system. **A**, Ctx–MMP-2; **B**, P75 – MMP-2; **C**, P76 – MMP-2; **D**, P77 – MMP-2; **E**, P78 – MMP-2.

**S2 Fig.Time course of RMSDs of Cα-atoms during MD simulations of peptide – MMP-2 complexes using the re-ranked top HPEPDOCK docking poses.** MMP-2, black; peptide, red.

**S3 Fig. MD simulations of the peptide – MMP-2 complexes using the using best scored docking pose obtained by the HADDOCK method.** Configurational entropy (black) and overlap of sampled region of subspace (red) of system. **A**, Ctx – MMP-2; **B**, P75 – MMP-2; **C**, P76 – MMP-2; **B**, P77 – MMP-2; **E**, P78 – MMP-2.

**S4 Fig. Time course of RMSDs of Cα-atoms during MD simulations of peptide – MMP-2 complexes using the using best scored docking pose obtained by the HADDOCK method.** MMP-2, black; peptide, red.

**S5 Fig. MD simulations of the peptide – MMP-2 complexes using the peptide – MMP-2 complexes obtained by AlphaFold2 docking.** Configurational entropy (black) and overlap of sampled region of subspace (red) of system. **A**, Ctx – MMP-2; **B**, P75 – MMP-2; **C**, P76 – MMP-2; **D**, P77 – MMP-2; **E**, P78 – MMP-2. Captured during 500 ns MD simulation.

**S6 Fig. Time course of RMSDs of Cα-atoms during MD simulations of peptide – MMP-2 complexes obtained by AlphaFold2 docking.** MMP-2, black; peptide, red.

**S7 Fig. SPR analysis of NRP-1 binding to P75 and P78.** NRP-1 was immobilized on an NTA chip, and peptides were injected over the chip surface. **(A)**Binding analysis of P75 to NRP-1, shown as the representative isotherm plot (left) and sensorgram (right). **(B)**Binding analysis of P78 to NRP-1, with the isotherm plot (left) and sensorgram (right). Both P75 and P78 exhibit weak binding to NRP-1. n=3, 4 replicates.

**S8 Fig. Representative Images of Wound-Healing U-87MG cells with Ctx and its C-terminal Fragments**.**A**, Control; **B**, Ctx; **C**, P75; **D**, P78; Left, 0 Hour; Right, 24 Hour. Percentage of wound closure was measured using the Wound_Healing_Size_tool_plugin in ImageJ.

**MD simulation trajectories, MD topology files, and pdb files from which figures are generated are available at** https://doi.org/10.5281/zenodo.17382049.

**XLXS file with raw data**

## References

1. Louis DN, Perry A, Wesseling P, Brat DJ, Cree IA, Figarella-Branger D, et al. The 2021 WHO Classification of Tumors of the Central Nervous System: a summary. Neuro-Oncol. 2021 Aug 2;23(8):1231–51.

2. Quesnel A, Karagiannis GS, Filippou PS. Extracellular proteolysis in glioblastoma progression and therapeutics. Biochim Biophys Acta BBA - Rev Cancer. 2020 Dec;1874(2):188428.

3. Rempe RG, Hartz AM, Bauer B. Matrix metalloproteinases in the brain and blood–brain barrier: Versatile breakers and makers. J Cereb Blood Flow Metab. 2016 Sep;36(9):1481–507.

4. Cui N, Hu M, Khalil RA. Biochemical and Biological Attributes of Matrix Metalloproteinases. Prog Mol Biol Transl Sci. 2017;147:1–73.

5. Visse R, Nagase H. Matrix metalloproteinases and tissue inhibitors of metalloproteinases: structure, function, and biochemistry. Circ Res. 2003;92(8):827–39.

6. Nagase H, Visse R, Murphy G. Structure and function of matrix metalloproteinases and TIMPs. Cardiovasc Res. 2006;69(3):562–73.

7. Yong VW, Power C, Forsyth P, Edwards DR. Metalloproteinases in biology and pathology of the nervous system. Nat Rev Neurosci. 2001 Jul;2(7):502–11.

8. Coussens LM, Fingleton B, Matrisian LM. Matrix metalloproteinase inhibitors and cancer—trials and tribulations. Science. 2002;295(5564):2387–92.

9. Forsyth PA, Wong H, Laing TD, Rewcastle NB, Morris DG, Muzik H, et al. Gelatinase-A (MMP-2), gelatinase-B (MMP-9) and membrane type matrix metalloproteinase-1 (MT1-MMP) are involved in different aspects of the pathophysiology of malignant gliomas. Br J Cancer. 1999 Apr;79(11/12):1828–35.

10. Voit-Ostricki L, Lovas S, Watts CR. Conformation and Domain Movement Analysis of Human Matrix Metalloproteinase-2: Role of Associated Zn2+ and Ca2+ Ions. Int J Mol Sci. 2019 Aug 27;20(17):4194.

11. Fang J, Shing Y, Wiederschain D, Yan L, Butterfield C, Jackson G, et al. Matrix metalloproteinase-2 is required for the switch to the angiogenic phenotype in a tumor model. Proc Natl Acad Sci. 2000;97(8):3884–9.

12. Du R, Petritsch C, Lu K, Liu P, Haller A, Ganss R, et al. Matrix metalloproteinase-2 regulates vascular patterning and growth affecting tumor cell survival and invasion in GBM. Neuro-Oncol. 2008;10(3):254–64.

13. Gong J, Zhu S, Zhang Y, Wang J. Interplay of VEGFa and MMP2 regulates invasion of glioblastoma. Tumor Biol. 2014 Dec 1;35(12):11879–85.

14. Ramachandran RK, Sørensen MD, Aaberg-Jessen C, Hermansen SK, Kristensen BW. Expression and prognostic impact of matrix metalloproteinase-2 (MMP-2) in astrocytomas. Ulasov I, editor. PLOS ONE. 2017 Feb 24;12(2):e0172234.

15. Jäälinojä J, Herva R, Korpela M, Höyhtyä M, Turpeenniemi-Hujanen T. Matrix metalloproteinase 2 (MMP-2) immunoreactive protein is associated with poor grade and survival in brain neoplasms. J Neurooncol. 2000;46(1):81–90.

16. Han L, Sheng B, Zeng Q, Yao W, Jiang Q. Correlation between MMP2 expression in lung cancer tissues and clinical parameters: a retrospective clinical analysis. BMC Pulm Med. 2020 Oct 28;20(1):283.

17. Tonn JC, Kerkau S, Hanke A, Bouterfa H, Mueller JG, Wagner S, et al. Effect of synthetic matrix-metalloproteinase inhibitors on invasive capacity and proliferation of human malignant gliomas In vitro. Int J Cancer. 1999 Mar 1;80(5).

18. Zhong Y, Lu YT, Sun Y, Shi ZH, Li NG, Tang YP, et al. Recent opportunities in matrix metalloproteinase inhibitor drug design for cancer. Expert Opin Drug Discov. 2018 Jan 2;13(1):75–87.

19. DeBin JA, Strichartz GR. Chloride channel inhibition by the venom of the scorpion Leiurus quinquestriatus. Toxicon. 1991 Jan;29(11):1403–8.

20. DeBin JA, Maggio JE, Strichartz GR. Purification and characterization of chlorotoxin, a chloride channel ligand from the venom of the scorpion. Am J Physiol-Cell Physiol. 1993 Feb 1;264(2):C361–9.

21. Ojeda PG, Chan LY, Poth AG, Wang CK, Craik DJ. The role of disulfide bonds in structure and activity of chlorotoxin. Future Med Chem. 2014 Oct 1;6(15):1617–28.

22. Gregory A, Voit-Ostricki L, Lovas S, Watts C. Effects of Selective Substitution of Cysteine Residues on the Conformational Properties of Chlorotoxin Explored by Molecular Dynamics Simulations. Int J Mol Sci. 2019 Mar 13;20(6):1261.

23. Olsen ML, Schade S, Lyons SA, Amaral MD, Sontheimer H. Expression of voltage-gated chloride channels in human glioma cells. J Neurosci Off J Soc Neurosci. 2003 Jul 2;23(13):5572–82.

24. Soroceanu L, Gillespie Y, Khazaeli MB, Sontheimer H. Use of chlorotoxin for targeting of primary brain tumors. Cancer Res. 1998 Nov 1;58(21):4871–9.

25. Lyons SA, O’Neal J, Sontheimer H. Chlorotoxin, a scorpion-derived peptide, specifically binds to gliomas and tumors of neuroectodermal origin. Glia. 2002 Aug 1;39(2):162–73.

26. Ullrich N, Bordey A, Gillespie GY, Sontheimer H. Expression of voltage-activated chloride currents in acute slices of human gliomas. Neuroscience. 1998 Jan;83(4):1161–73.

27. Maertens C, Wei L, Tytgat J, Droogmans G, Nilius B. Chlorotoxin does not inhibit volume-regulated, calcium-activated and cyclic AMP-activated chloride channels. Br J Pharmacol. 2000 Feb;129(4):791–801.

28. Deshane J, Garner CC, Sontheimer H. Chlorotoxin Inhibits Glioma Cell Invasion via Matrix Metalloproteinase-2. J Biol Chem. 2003 Feb 7;278(6):4135–44.

29. Costa PM, Cardoso AL, Mendonça LS, Serani A, Custódia C, Conceição M, et al. Tumor-targeted chlorotoxin-coupled nanoparticles for nucleic acid delivery to glioblastoma cells: a promising system for glioblastoma treatment. Mol Ther-Nucleic Acids. 2013 Jun 18;2(E100).

30. Maertens C. Chlorotoxin does not inhibit volume-regulated, calcium-activated and cyclic AMP-activated chloride channels.

31. Qin C, He B, Dai W, Lin Z, Zhang H, Wang X, et al. The impact of a chlorotoxin-modified liposome system on receptor MMP-2 and the receptor-associated protein ClC-3. Biomaterials. 2014 Jul 1;35(22):5908–20.

32. John A. DeBin. Purification and Characterization of Chlorotoxin, a Chloride Channel Ligand from the Venom of Scorpion.

33. Othman H, Wieninger SA, ElAyeb M, Nilges M, Srairi-Abid N. In Silico prediction of the molecular basis of ClTx and AaCTx interaction with matrix metalloproteinase-2 (MMP-2) to inhibit glioma cell invasion. J Biomol Struct Dyn. 2017 Oct 3;35(13):2815–29.

34. Alam M, Ali SA, Abbasi A, Kalbacher H, Voelter W. Design and Synthesis of a Peptidyl-FRET Substrate for Tumor Marker Enzyme human Matrix Metalloprotease-2 (hMMP-2). Int J Pept Res Ther. 2012 Sep 1;18(3):207–15.

35. Farkas S, Cioca D, Murányi J, Hornyák P, Brunyánszki A, Szekér P, et al. Chlorotoxin binds to both matrix metalloproteinase 2 and neuropilin 1. J Biol Chem. 2023 Sep;299(9):104998.

36. Cedars-Sinai Medical Center (Responsible Party). Fluorescence Detection of Adult Primary Central Nervous System Tumors With Tozuleristide and the Canvas System. Available from: https://clinicaltrials.gov/study/NCT04743310

37. Yamada M, Miller DM, Lowe M, Rowe C, Wood D, Soyer HP, et al. A first-in-human study of BLZ-100 (tozuleristide) demonstrates tolerability and safety in skin cancer patients. Contemp Clin Trials Commun. 2021 Sep;23:100830.

38. Blaze Bioscience Australia Pty Ltd. Safety Study of a Fluorescent Marker to Visualize Cancer Cells. Available from: https://clinicaltrials.gov/study/NCT02097875

39. Patil CG, Walker DG, Miller DM, Butte P, Morrison B, Kittle DS, et al. Phase 1 Safety, Pharmacokinetics, and Fluorescence Imaging Study of Tozuleristide (BLZ-100) in Adults With Newly Diagnosed or Recurrent Gliomas. Neurosurgery. 2019 Oct;85(4):E641–9.

40. Kobets AJ, Nauen D, Lee A, Cohen AR. Unexpected Binding of Tozuleristide “Tumor Paint” to Cerebral Vascular Malformations: A Potentially Novel Application of Fluorescence-Guided Surgery. Neurosurgery. 2021 Aug;89(2):204–11.

41. City of Hope Medical Center. Chimeric Antigen Receptor (CAR) T Cells With a Chlorotoxin Tumor-Targeting Domain for the Treatment of MMP2+ Recurrent or Progressive Glioblastoma. Available from: https://clinicaltrials.gov/study/NCT04214392

42. Wang D, Starr R, Chang WC, Aguilar B, Alizadeh D, Wright SL, et al. Chlorotoxin-directed CAR T cells for specific and effective targeting of glioblastoma. Sci Transl Med. 2020 Mar 4;12(533):eaaw2672.

43. Tudor T, Binder ZA, O’Rourke DM. CAR T Cells. Neurosurg Clin N Am. 2021 Apr;32(2):249–63.

44. Dastpeyman M, Giacomin P, Wilson D, Nolan MJ, Bansal PS, Daly NL. A C-Terminal Fragment of Chlorotoxin Retains Bioactivity and Inhibits Cell Migration. Front Pharmacol. 2019;10:250.

45. Berman HM, Westbrook J, Feng Z, Gilliland G, Bhat TN, Weissig H, et al. The Protein Data Bank. Nucleic Acids Res. 2000 Jan 1;28(1):235–42.

46. Lippens G, Najib J, Wodak SJ, Tartar A. NMR Sequential Assignments and Solution Structure of Chlorotoxin, a Small Scorpion Toxin That Blocks Chloride Channels. Biochemistry. 1995 Jan 10;34(1):13–21.

47. Tao H, Zhao X, Zhang K, Lin P, Huang SY. Docking cyclic peptides formed by a disulfide bond through a hierarchical strategy. Bioinformatics. 2022 Sep 2;38(17):4109–16.

48. Honorato RV, Koukos PI, Jiménez-García B, Tsaregorodtsev A, Verlato M, Giachetti A, et al. Structural Biology in the Clouds: The WeNMR-EOSC Ecosystem. Front Mol Biosci. 2021 Jul 28;8.

49. Jumper J, Evans R, Pritzel A, Green T, Figurnov M, Ronneberger O, et al. Highly accurate protein structure prediction with AlphaFold. Nature. 2021 Aug;596(7873):583–9.

50. Zhou P, Jin B, Li H, Huang SY. HPEPDOCK: a web server for blind peptide–protein docking based on a hierarchical algorithm. Nucleic Acids Res. 2018 Jul 2;46(W1):W443– 50.

51. Dominguez C, Boelens R, Bonvin AMJJ. HADDOCK: A Protein−Protein Docking Approach Based on Biochemical or Biophysical Information. J Am Chem Soc. 2003 Feb 19;125(7):1731–7.

52. Mirdita M, Schütze K, Moriwaki Y, Heo L, Ovchinnikov S, Steinegger M. ColabFold: making protein folding accessible to all. Nat Methods. 2022 Jun;19(6):679–82.

53. Krieger E, Vriend G. New ways to boost molecular dynamics simulations. J Comput Chem. 2015 May 15;36(13):996–1007.

54. Land H, Humble MS. YASARA: A Tool to Obtain Structural Guidance in Biocatalytic Investigations. Methods Mol Biol Clifton NJ. 2018;1685:43–67.

55. Wang LP, McKiernan KA, Gomes J, Beauchamp KA, Head-Gordon T, Rice JE, et al. Building a More Predictive Protein Force Field: A Systematic and Reproducible Route to AMBER-FB15. J Phys Chem B. 2017 Apr 27;121(16):4023–39.

56. Essmann U, Perera L, Berkowitz ML, Darden T, Lee H, Pedersen LG. A smooth particle mesh Ewald method. J Chem Phys. 1995 Nov 15;103(19):8577–93.

57. Abraham MJ, Murtola T, Schulz R, Páll S, Smith JC, Hess B, et al. GROMACS: High performance molecular simulations through multi-level parallelism from laptops to supercomputers. SoftwareX. 2015 Sep 1;1-2:19–25.

58. Huang J, Rauscher S, Nawrocki G, Ran T, Feig M, de Groot BL, et al. CHARMM36m: an improved force field for folded and intrinsically disordered proteins. Nat Methods. 2017 Jan;14(1):71–3.

59. Jorgensen WL, Chandrasekhar J, Madura JD, Impey RW, Klein ML. Comparison of simple potential functions for simulating liquid water. J Chem Phys. 1983 Jul 15;79(2):926– 35.

60. Bussi G, Donadio D, Parrinello M. Canonical sampling through velocity rescaling. J Chem Phys. 2007 Jan 7;126(1):014101.

61. Berendsen HJC, Postma JPM, Van Gunsteren WF, DiNola A, Haak JR. Molecular dynamics with coupling to an external bath. J Chem Phys. 1984 Oct 15;81(8):3684–90.

62. Berk H, Bekker H, Berendsen HJC, Fraaije JGEM. LINCS: A linear constraint solver for molecular simulations. J Comput Chem. 1997;18(12):1463–72.

63. Parrinello M, Rahman A. Polymorphic transitions in single crystals: A new molecular dynamics method. J Appl Phys. 1981 Dec 1;52(12):7182–90.

64. Grubmüller H, Heller H, Windemuth A, Schulten K. Generalized Verlet Algorithm for Efficient Molecular Dynamics Simulations with Long-range Interactions. Mol Simul. 1991 Mar;6(1–3):121–42.

65. Valdés-Tresanco MS, Valdés-Tresanco ME, Valiente PA, Moreno E. gmx_MMPBSA: A New Tool to Perform End-State Free Energy Calculations with GROMACS. J Chem Theory Comput. 2021 Oct 12;17(10):6281–91.

66. Valdés-Tresanco MS, Valdés-Tresanco ME, Valiente PA, Moreno E. Supporting Information gmx_MMPBSA: a new tool aid to perform end-state free energy calculations with GROMACS files.

67. Karplus M, Kushick JN. Method for estimating the configurational entropy of macromolecules. Macromolecules. 1981 Mar;14(2):325–32.

68. Andricioaei I, Karplus M. On the calculation of entropy from covariance matrices of the atomic fluctuations. J Chem Phys. 2001 Oct 8;115(14):6289–92.

69. Hess B. Convergence of sampling in protein simulations. Phys Rev E. 2002 Mar 1;65(3):031910.

70. Daura X, Gademann K, Bernhard J, Seebach D, Van Gunsteren WF, Mark AE. Peptide Folding: When Simulation Meets Experiment. Angew Chem Int Ed. 1999;38(1–2):236–40.

71. Kamayirese S, Maity S, Dieckman LM, Hansen LA, Lovas S. Optimizing Phosphopeptide Structures That Target 14-3-3ε in Cutaneous Squamous Cell Carcinoma. ACS Omega. 2024 Jan 16;9(2):2719–29.

72. Kamayirese S, Maity S, Hansen LA, Lovas S. The Development of CDC25A-Derived Phosphoseryl Peptides That Bind 14-3-3ε with High Affinities. Int J Mol Sci. 2024 Jan;25(9):4918.

73. Díaz N, Suárez D. Molecular dynamics simulations of the active matrix metalloproteinase-2: positioning of the N-terminal fragment and binding of a small peptide substrate. Proteins. 2008 Jul;72(1):50–61.

74. Kesavan K, Ratliff J, Johnson EW, Dahlberg W, Asara JM, Misra P, et al. Annexin A2 Is a Molecular Target for TM601, a Peptide with Tumor-targeting and Anti-angiogenic Effects. J Biol Chem. 2010 Feb 12;285(7):4366–74.

75. Sonkin D, Thomas A, Teicher BA. Cancer treatments: Past, present, and future. Cancer Genet. 2024 Aug;286-287:18–24.

